# Dual MMP-9/12 Inhibition with AZD1236 Confers Neurovascular Protection and Reduces Post-Stroke Pain in Experimental Stroke Models

**DOI:** 10.64898/2026.08.23.746523

**Authors:** Milena De Felice, Saurabh Jain, Steven Reynolds, Tushar K Ghosh, Raymond Wong, Catherine B Lawrence, Matthew Worsley, Jilian Newton, Philip M Bath, Alistair Buchan, Iain Gardner, Arshad Majid

**Affiliations:** School of Clinical Dentistry, University of Sheffield, UK; The Neuroscience Institute, University of Sheffield, UK; Theralndx Lifesciences, India; School of Medicine and Population Health University of Sheffield, UK; Sheffield Institute for Translational Neuroscience, UK; Geoffrey Jefferson Brain Research Centre, Manchester Academic Health Science Centre, Northern Care Alliance NHS Foundation Trust, The University of Manchester, UK; Division of Neuroscience, Faculty of Biology, Medicine and Health, The University of Manchester, UK; Biomolecular Science Research Centre, Sheffield Hallam University, UK; Stroke Trials Unit, University of Nottingham, Nottingham, UK; University of Oxford, Radcliffe Department of Medicine, UK; Certara, Sheffield UK

**Keywords:** Stroke, MMP-9, MMP-12, in vivo, cerebroprotection, post-stroke pain

## Abstract

**Background:** Stroke remains a leading cause of death and disability worldwide. Matrix metalloproteinases (MMPs), particularly MMP-9 and MMP-12, contribute to early blood-brain barrier (BBB) disruption, neuroinflammation, haemorrhagic transformation, and intracerebral haemorrhage (ICH). Intravenous thrombolysis is the only widely used pharmacological therapy for acute ischaemic stroke, but its utility is limited by narrow eligibility criteria and haemorrhagic risk. Inhibition of MMPs in the acute phase may offer a complementary neurovascular protective strategy.

**Methods:** AZD1236, a selective dual MMP-9/-12 inhibitor, was evaluated in transient and permanent middle cerebral artery occlusion models and in a collagenase-induced ICH model in young, aged, obese, and female mice. Drug or vehicle was administered 2-6 hours after stroke onset. Outcomes included infarct or haematoma volume, BBB integrity, neurological function, and pain related behaviours.

**Results:** AZD1236 given within 2-4 hours after ischaemic or haemorrhagic insult significantly reduced infarct and haematoma volumes, improved short-and long-term neurological scores, and preserved BBB integrity, whereas treatment at 6 hours was largely ineffective. AZD1236 also attenuated the development of post-stroke mechanical allodynia and thermal hyperalgesia. Mechanistically, treatment reduced MMP-9 and MMP-12 activity, increased tight junction protein expression, and dampened inflammatory responses.

**Conclusions:** Dual inhibition of MMP-9/-12 with AZD1236 confers robust neurovascular protection and mitigates post-stroke pain across clinically relevant models of ischaemic and haemorrhagic stroke. These findings provide a strong preclinical rationale for clinical evaluation of dual MMP-9/12 inhibition as an adjunctive neuroprotective strategy for acute stroke.

## INTRODUCTION

Stroke is the second leading cause of death and a major cause of long-term disability worldwide (1–5). Despite advances in reperfusion therapy, treatment options remain limited. Intravenous thrombolysis with alteplase or tenecteplase remains the main pharmacological approach for acute ischaemic stroke, but its use is constrained by a narrow treatment window and stringent eligibility criteria (5–13). Although advanced imaging has enabled extension of treatment windows, only a minority of patients ultimately receive thrombolysis. Endovascular thrombectomy has transformed outcomes in selected patients with large-vessel occlusion and can be effective up to 24 hours from onset (6, 7). However, many patients remain ineligible for either therapy because of clot location, comorbidities, delayed presentation, or logistical barriers, and thus receive no effective acute intervention (14–16). There is therefore an urgent need for adjunctive neuroprotective strategies that can be combined with reperfusion and extend benefit to patients who are not candidates for thrombolysis or thrombectomy. Reperfusion does not address the cascade of secondary injury triggered by ischaemia. Excitotoxicity, oxidative stress, neuroinflammation, and disruption of the blood-brain barrier (BBB) contribute to infarct expansion, haemorrhagic transformation, and poor functional outcomes (17–24). A neuroprotective therapy that preserves the neurovascular unit, limits BBB breakdown, and is compatible with existing reperfusion approaches would represent a major advance in stroke care.

Among the mediators of secondary injury (17, 18, 20, 23), matrix metalloproteinases (MMPs) play a central role. MMP-9 is rapidly upregulated in endothelial cells after the onset of ischaemia, drives BBB disruption, and increases the risk of haemorrhagic transformation, particularly after thrombolysis (25–37). MMP-12, expressed predominantly by infiltrating macrophages and microglia, degrades extracellular matrix and myelin proteins and can activate pro-MMP-9, thereby amplifying tissue damage (30, 33, 36, 38–41). Inhibition of either enzyme reduces injury in experimental stroke models, but dual inhibition may provide broader and more sustained neurovascular protection than targeting either alone (30, 42–44).

AZD1236 is a selective, orally bioavailable inhibitor of human MMP-9 and MMP-12, with nanomolar potency and >350-fold selectivity over other MMP family members. Originally developed for chronic obstructive pulmonary disease, AZD1236 has demonstrated acceptable safety and tolerability in Phase I and II clinical trials (45, 46). The extent of AZD1236 penetration across the intact blood-brain barrier has not been fully characterised. However, because BBB disruption occurs rapidly following cerebral ischaemia, systemically administered AZD1236 may gain greater access to injured brain tissue during the acute phase of stroke. In preclinical models of spinal cord injury, AZD1236 reduced oedema and improved functional outcomes (47), supporting its translational potential in acute neurovascular injury.

Beyond the acute phase, stroke survivors frequently experience central post-stroke pain (CPSP), a debilitating neuropathic complication linked to persistent neuroinflammation and microglial activation (48–58). MMP-9 has been implicated as an initiator of neuropathic pain through interleukin-1β cleavage and microglial activation, and MMP-9/-12 inhibition attenuates pain hypersensitivity in spinal cord injury models (47, 59–61), suggesting that dual inhibition may also reduce CPSP. Here, AZD1236 was systematically evaluated in transient andpermanent models of ischaemic stroke in young, aged, obese, male and female mice, in line with STAIR recommendations (53). Furthermore, AZD1236 was tested in a collagenase-induced intracerebral haemorrhage model to determine whether dual MMP-9/12 inhibition confers broader neurovascular protection in haemorrhagic stroke.

### Materials and Methods Animals

Young male (6–8 weeks, 20–25 g), old male (70–74 weeks, 40–45 g), young female (23–25 weeks, 20–25 g), and old female (70–74 weeks, 35–40 g) C57BL/6J mice were purchased from Charles River (UK). Animals were housed under standard conditions (12-hour light/dark cycle, food and water *ad libitum*, 5 mice per cage).

### Obese Mice

Diet-induced obesity was generated in male 6-8 weeks old C57BL/6J mice by feeding a high-fat diet (60% kcal from fat; Test Diets®) for 4 months prior to stroke surgery, as previously described (62, 63).

All procedures were carried out under a licence issued by the UK Home Office in accordance with the Animals (Scientific Procedures) Act, 1986, and were approved by the local ethics committee (70/8408 and PP2294383 and PP9466981).

### Randomisation, Blinding, and Sample Size

Animals were randomly assigned to treatment or vehicle groups using a computer-generated random number function in Excel. All surgical procedures, drug administrations, and outcome assessments were performed by investigators blinded to treatment allocation; animals were labelled with anonymised codes, and treatment identity was revealed only after data analysis was finalised. Sample size calculations were performed using published data (51) in-house pilot studies, and feasibility considerations for each model. Final group sizes for each experiment are reported in the corresponding figure legends. For infarct volume, group sizes were estimated to detect a 20% difference with 80% power at α = 0.05. For Garcia neurological scores, group sizes were estimated to detect an absolute between-group difference of approximately 1.5 points with 80% power at α = 0.05.

### Drug Administration

AZD1236 was dissolved in 40% 2-hydroxypropyl-β-cyclodextrin in distilled water. Vehicle-treated animals received the same solution without drug. The initial 200 mg/kg dose was administered as a 100 mg/kg intravenous bolus immediately followed by 100 mg/kg oral gavage. Subsequent doses of 200 mg/kg were given orally starting 24 hours after stroke and continued once daily for 6 days.

### Justification for chosen dose

AZD1236 was administered at 200 mg/kg to achieve high systemic exposure and near-maximal inhibition of MMP-9 and MMP-12, based on prior preclinical data demonstrating low-nanomolar potency against both murine enzymes and dose-dependent pharmacologic activity in rodent models (47, 64). This regimen was further supported by pharmacokinetic and pharmacodynamic data from studies of AZD1236 in spinal cord injury where sustained target engagement in the CNS was associated with functional and histological benefit (47).

### Transient Middle Cerebral Artery Occlusion (tMCAO)

Mice were anaesthetised with 5% isoflurane and maintained at 1.5% isoflurane in oxygen-enriched air. Body temperature was maintained at 37 ± 0.5 °C using a feedback-controlled heating pad. The left common carotid artery (CCA) was ligated, the external carotid artery (ECA) tied, and a silicone-coated monofilament (Doccol Corporation, USA) was advanced into the internal carotid artery (ICA) to occlude the middle cerebral artery (MCA). The occlusion time was 30 minutes for obese mice, 45 minutes for young and aged female and aged male mice, and 60 minutes for young male mice, to compensate for known sex-and age-dependent differences in ischaemic sensitivity and infarct evolution and to achieve comparable cortical infarct volumes across groups while maintaining acceptable mortality and neurological severity (65). Reperfusion was confirmed by an increase in cerebral blood flow on laser Doppler flowmetry (51).

### Permanent Middle Cerebral Artery Occlusion (pMCAO)

Mice were anaesthetised with 5% isoflurane and maintained at 1.5% isoflurane. Body temperature was maintained at 37 ± 0.5 °C. A small craniotomy was performed to expose the MCA, which was occluded proximally and distally using electrocoagulation (66). The wound was sutured and mice recovered in a warming box (32 °C) for 30 minutes before return to the home cages.

### Haemorrhagic Stroke Model

Intracerebral haemorrhage was induced by stereotaxic injection of collagenase VII. A burr hole was drilled at stereotaxic coordinates 2 mm lateral and 1 mm anterior to bregma, depth 3.5 mm. Collagenase VII (0.075 U in 0.5 μL saline) or saline control was injected into the striatum over 2 minutes using a Hamilton syringe. The needle was withdrawn slowly after 10 minutes and the wound sutured (67).

### Infarct and Haemorrhage Volume Analysis

At 48 hours post-stroke, brains were removed, sliced into 1 mm coronal sections, and incubated in 2% TTC at 37 °C for 20 minutes. Infarct areas were measured with ImageJ, corrected for oedema, and volumes calculated using the formula: [1 – (ipsilateral hemisphere – infarct volume) / contralateral hemisphere] × 100 (51).

### Neurological Assessment

Neurological deficits were assessed using the Garcia 18-point scale before surgery and at predefined intervals post-stroke. The scale evaluates spontaneous activity, limb symmetry, forelimb extension, climbing, proprioception, grip strength, and vibrissae response; lower scores indicate greater deficit (68, 69). Assessors were blinded to treatment group.

### Pain Behaviours

Baseline mechanical and thermal sensitivity were measured prior to surgery. Mechanical allodynia was assessed using a 2 g von Frey filament (10 applications; withdrawal responses recorded) (70). Thermal hyperalgesia was assessed with the Hargreaves test (baseline latencies 7–9 seconds; 10-second cut-off) (70). Testing was repeated on days 7, 14, 21, and 28 after ischaemia onset. Animals were monitored to confirm intact withdrawal reflexes prior to testing, ensuring that behavioural responses were not confounded by gross motor deficits.

### Blood-Brain Barrier Integrity

For Evans Blue extravasation, mice received an intravenous bolus of 2% Evans Blue at 48 hours post-MCAO. After 45 minutes, brains were extracted, weighed, and incubated in formamide at 60 °C for 72 hours, and dye content was quantified spectrophotometrically at 620 nm (71). For MRI, at 24 hours post-tMCAO mice underwent scanning on a 7T Bruker Avance III MRI scanner (Bruker BioSpin GmbH & Co, Ettlinggen, Germany), equipped with 660 mT/m gradients, 86mm ID volume transmit resonator coil and 4-phase mouse brain receive coil. During scanning mice were anaesthetised using 1-2% isoflurane and enriched oxygen, with body temperature maintained through a combination of heated water and electrical blanket. The animal movement was restricted with ear bars, nose cone and bite bar. Breathing and body temperature were monitored. Images were acquired using a RARE-VTR sequence (slice thickness 0.75 mm; in-plane resolution 0.25 × 0.25 mm; FOV 20×20 mm; 5 TRs 1000, 1568, 2364, 3707, 10000 ms; 4 TEs 13, 39, 65 91ms; RARE factor 2), before and after intraperitoneal gadolinium administration (1:4 DOTAREM in saline, 100 μL). The post-contrast RARE-VTR sequence was started approximately 30 minutes after injection. T1 and T2 maps were generated using custom MATLAB scripts (MathWorks, Natick, MA) to fit T1 saturation recovery and T2 decay equations respectively using a non-linear least squares method. Regions of interest (ROIs) were manually delineated for all image slices containing the stroke/infarct region. For comparison a contra-lateral ROI was created by reflecting the infarct ROI about the mid-line of the brain by an investigator blinded to treatment. Mean pre-to post-gadolinium T1 signal ratios were calculated for each ROI and used as an index of BBB permeability

### Western Blotting

Proteins were extracted from ischaemic brain tissue at day 3 post-tMCAO. Lysates (25 μg) were separated on Tris–Acetate gels, transferred to PVDF membranes, and probed with antibodies against MMP-9 (1:500, Santa Cruz, sc-393859), MMP-12 (1:500, Santa Cruz, sc-390863), claudin-5 (1:1000, ThermoFisher, 4C3C2), and ZO-1 (1:1000, ThermoFisher, 61-7300). HRP-conjugated secondary antibodies (Abcam ab6721, ab205719) were used, and bands were detected by enhanced chemiluminescence. Band intensities were quantified using ImageJ and normalised to housekeeping proteins (α-tubulin, Abcam; β-actin, ThermoFisher) (30).

### Gelatinase inhibitor activity assay

Plasma and brain gelatinase inhibitory activity were assessed using a fluorogenic DQ-gelatin assay (EnzChek Gelatinase/Collagenase Assay Kit, Thermo Fisher Scientific, E12055). At 24 hours post-pMCAO, mice were euthanised, brains were rapidly dissected and snap frozen, and blood was collected into heparinised tubes and centrifuged at 2,000 × g for 10 minutes at 4 °C to obtain plasma. Brain and plasma samples were aliquoted and stored at −80 °C until analysis. Prior to the assay, ipsilateral cortex/striatum was homogenised in 50 mM Tris-HCl (pH 7.4) containing 10 mM CaCl₂ and 0.1% Triton X-100, centrifuged (10,000 × g, 10 minutes), and supernatants collected. For the inhibitory assay, a fixed amount of active gelatinase was pre-incubated with diluted plasma or brain samples (final dilution 1:10 in reaction buffer) for 15 minutes at 37 °C prior to substrate addition. DQ-gelatin substrate was then added in 1× reaction buffer to a final volume of 100 μL per well in black-walled, clear-bottom 96-well plates. Fluorescence was measured kinetically at 37 °C using a microplate reader (excitation 495 nm, emission 515 nm) every 5 minutes for up to 120 minutes. This assay reflects the net inhibitory capacity of the biological sample against a defined gelatinase substrate rather than direct measurement of endogenous MMP activity. Gelatinase activity was quantified as the linear rate of fluorescence increase (ΔRFU/min) after subtraction of background fluorescence from substrate-only wells. Inhibitory activity was expressed as percentage reduction in enzyme activity relative to enzyme-only controls. All samples were assayed in triplicate.

### Pharmacokinetics

At terminal anaesthesia (pentobarbital, 200 mg/kg intraperitoneally), blood was collected by cardiac puncture in mice with tMCAO. Plasma was separated (11,000 rpm, 5 minutes, 4 °C) and stored at −80 °C. AZD1236 concentrations were quantified by LC–MS/MS at Sheffield Hallam University using an Agilent Infinity II 1290 HPLC system and an XBridge BEH Amide column. Frozen plasma and brain samples were thawed at room temperature. Plasma samples were then vortex mixed and centrifuged at 2400 g for 5 minutes and 25 µl of supernatant was used for analysis, whereas brain samples were homogenised using a tissue grinder and 100mg of homogenate was used for analysis. Each aliquot of plasma or brain sample was mixed with 150 µl of internal standard (500 nM AZD3342 in 0.2% formic acid in acetonitrile) in a glass chromatography vial. Vials were vortex mixed for 30 s and then centrifuged at 2500 g for 10 minutes. 50 µl of supernatant was removed and mixed with 300 µl of sterile ultrapure water and vortexed for 30 s. 10 µl was injected and analysed in a multiple reaction monitoring (MRM) assay, using liquid chromatography (LC) coupled tandem mass spectrometry (MS/MS), with an electrospray ionisation (ESI) source (LC-ESI-MRM-MS/MS).

Data acquisition and analysis were achieved using MassHunter Acquisition and Qualitative Analysis version B.0.10.0. The following ion transitions were used for AZD1236 (m/z 416.2 to 197.1) and for the internal standard AZD3342 (m/z 402.2 - 190.2).

### Statistical Analysis

Data are presented as mean ± SEM. Normality was assessed using the Shapiro-Wilk test, as previously described in rodent models of tMCAO and pMCAO (72, 73). For comparisons between two independent groups, two-tailed unpaired Welch’s t-test were used. For therapeutic-window experiments (2,4, and 6 hours), each AZD1236-treated group was compared with its contemporaneous vehicle control. Experiments including a single vehicle group and multiple treatment groups were analysed using one-way ANOVA followed by Dunnett’s multiple comparisons test. Longitudinal Garcia neurological score and pain-related behaviours were analysed using two-way repeated-measures ANOVA or mixed-effects model, as appropriate, followed by multiple-comparisons testing. MRI data were analysed in MATLAB using Kruskal-Wallis followed by Bonferroni-adjusted post hoc comparisons. Analyses were performed using GraphPad Prism v9 software. Sample sizes represent independent biological replicates as indicated in the corresponding figure legends. Statistical significance was defined as p < 0.05.

## Results

### Pharmacokinetics

Administration of AZD1236 (200 mg/kg) resulted in substantial plasma and brain concentrations in both young and aged mice of both sexes (Fig. 1a,b). AZD1236 exhibits substantial plasma and brain protein binding in mice (93.8% and 94.0%, respectively), corresponding to free fractions of 0.062 and 0.0597. After correction for binding, the expected brain:plasma ratio approaches unity under conditions of free diffusion, although observed brain concentrations were approximately 50% of plasma levels at 6 hours post-dose, suggesting partial restriction of CNS penetration.

**Figure 1.**
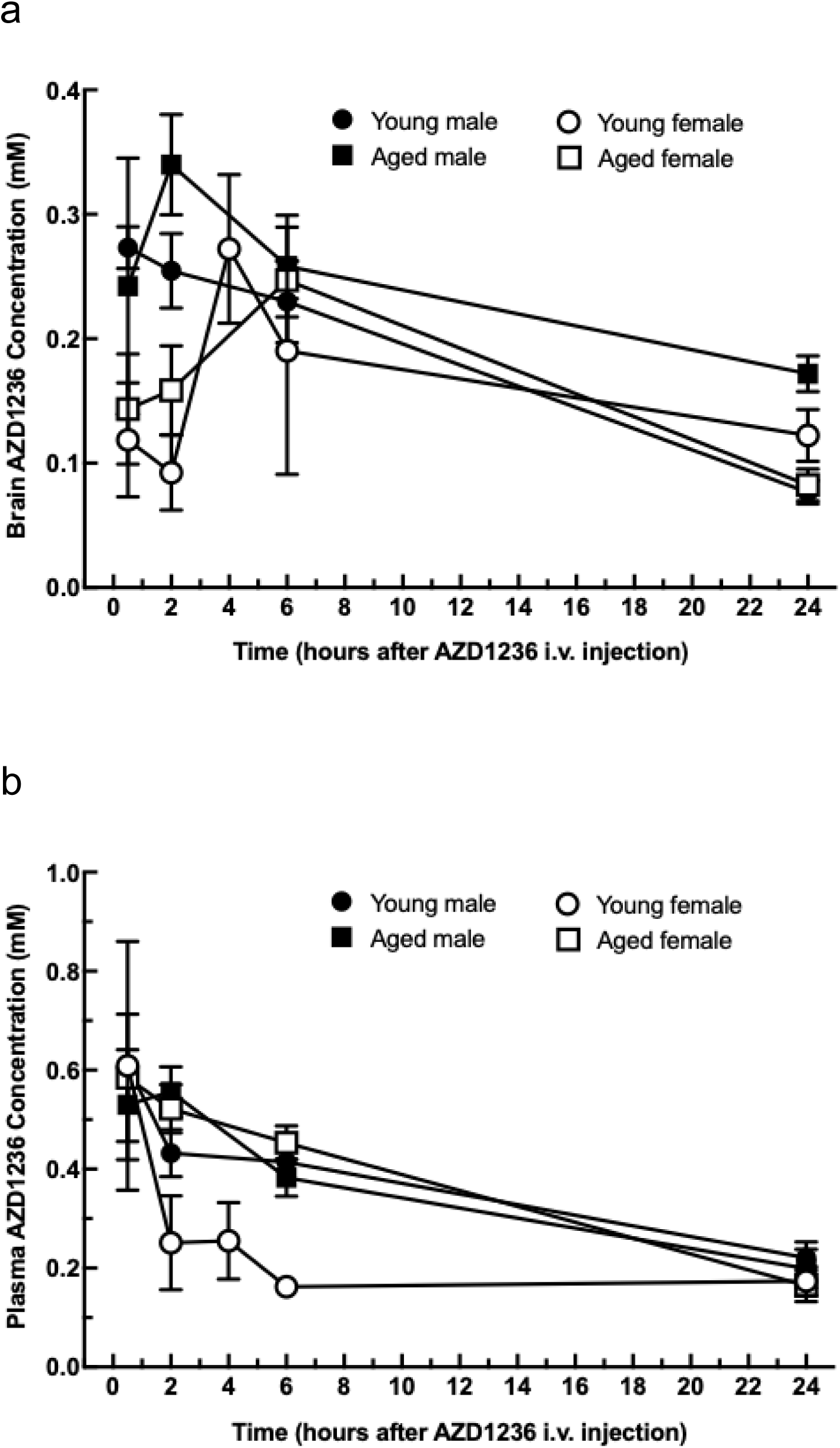
AZD1236 achieves robust plasma and brain exposure after tMCAO. (a) Brain and (b) plasma concentrations of AZD1236 in young and aged male and female mice after transient middle cerebral artery occlusion. AZD1236 (200 mg/kg) was administered 2 h after ischaemia onset and levels measured over 24 h, demonstrating sustained plasma and brain exposure across ages and sexes. Data are mean ± SEM, n = 5-6 per group.*p < 0.05 versus vehicle.

At 200 mg/kg, total brain concentrations ranged from 0.1–0.3 mM and plasma concentrations from 0.2–0.6 mM. This corresponds to estimated free plasma concentrations of 12.4–37.2 µM and free brain concentrations of 6.0–17.9 µM. These exposures substantially exceed the murine IC50 values for both targets (MMP-9: 270 nM; MMP-12: 140 nM), achieving approximately 46–137-fold and 89–265-fold target coverage in plasma, and 22–66-fold and 43–128-fold coverage in brain, respectively. These values reflect a substantial species difference in potency. AZD1236 was optimised against the human enzymes (IC50 4.5 nM for MMP-9 and 6.1 nM for MMP-12), with activity at rodent orthologues approximately 20-to 50-fold lower; the murine values used here correspond to 60-fold and 23-fold reductions respectively, consistent with that differential. The 200 mg/kg regimen therefore reflects the exposure required to cover the comparatively insensitive murine enzymes rather than an intrinsically high dose requirement.

Together, these data indicate that the selected dosing regimen achieves robust systemic and CNS pharmacologically active exposure, sufficient to ensure near-complete inhibition of MMP-9 and MMP-12 activity in both compartments, consistent with previously reported exposure–response relationships in CNS injury paradigms. Supplementary Fig. 10 represents PK data in obese mice dosed at 100mg/kg, whereas Fig 1 shows young and aged mice dosed at 200mg/kg. These datasets are therefore not directly comparable and are presented to demonstrate exposure within each experimental setting rather than cross-cohort equivalence. Furthermore, the objective of this study was to establish whether sustained dual MMP-9/12 inhibition is sufficient to confer neurovascular protection rather than to define a clinically translatable dosing regimen, hence the dosing regimen chosen.

### Efficacy in ischaemic stroke models

In the pMCAO model, AZD1236 administered 2 or 4 hours post-occlusion in young male mice, significantly reduced infarct volume at 48 hours compared with vehicle controls (47% and 34% reductions; p < 0.01 and p < 0.05, respectively; Fig. 2a). Administration at 6 hours did not reduce infarct size. Longitudinal assessment of neurological function using the Garcia scores, demonstrated a significant effect of treatment over time. AZD1236 administration at 2 or 4 hours also improved long-term functional recovery compared with vehicle controls, with significantly higher Garcia neurological scores maintained throughout the 28 days observation period (p < 0.01) whereas no benefit was observed when dosing was initiated at 6 hours (Fig. 2b).

**Figure 2.**
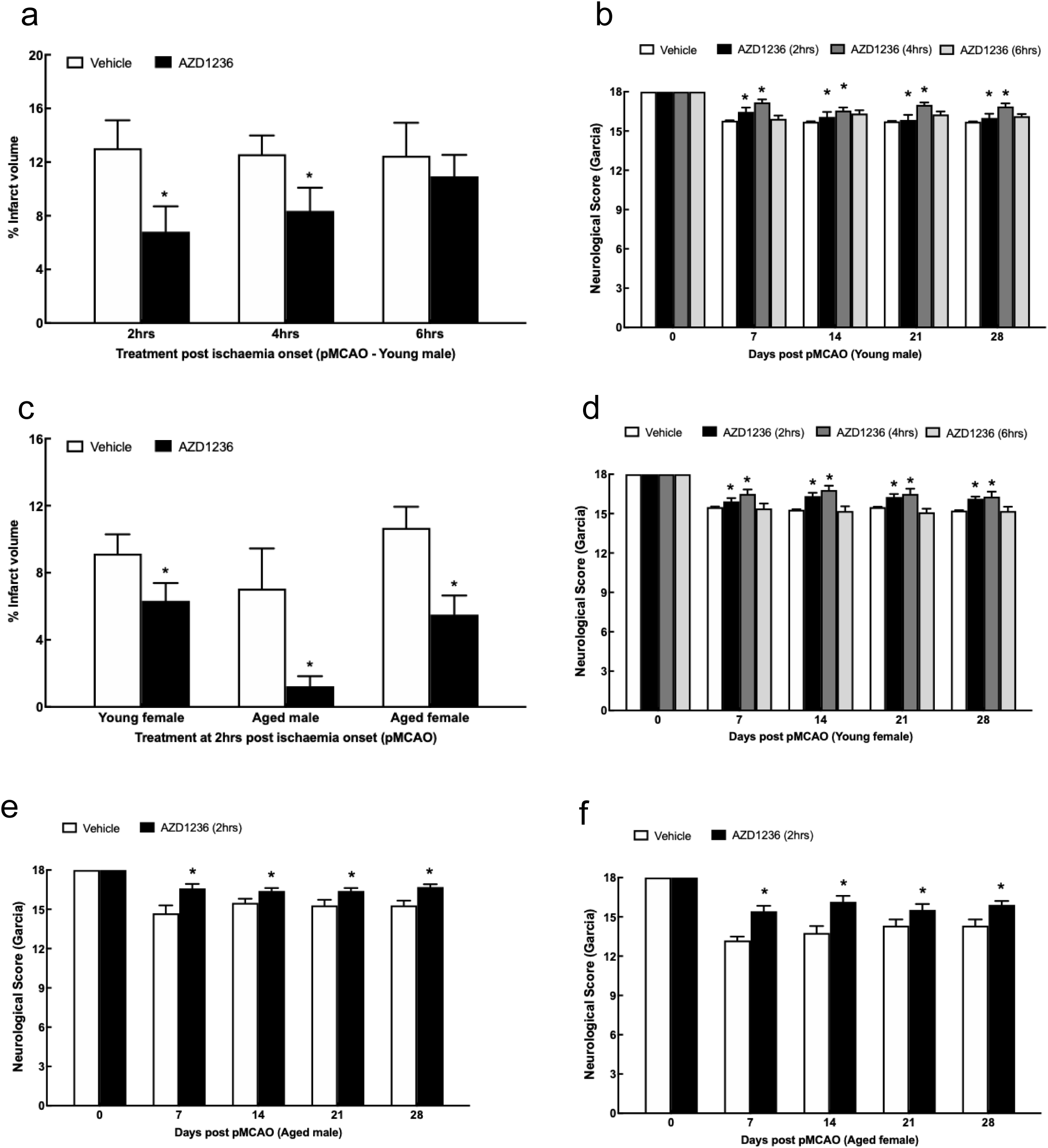
Time-dependent neuroprotection by AZD1236 following pMCAO. Infarct volume in young (a,c) and aged (e) mice, and longitudinal neurological function assessed using the 18-point Garcia score in young (b,d) and aged (f) mice following pMCAO. AZD1236 (200 mg/kg) or vehicle was administered 2, 4 or 6 h after ischaemia onset. Infarct-volume data are presented as mean ± SEM (n = 6–8 biological replicates per group) and were analysed using two-tailed unpaired Welch’s t tests comparing each AZD1236-treated group with its contemporaneous vehicle control. Garcia scores are presented as mean ± SEM (n = 10–15 biological replicates per group) and were analysed using a mixed-effects model (or two-way repeated-measures ANOVA where appropriate) followed by multiple-comparisons testing. *P < 0.05 versus the corresponding vehicle group.

The protective profile was replicated in young female mice, where AZD1236 (200 mg/kg, 2–4 hours) reduced infarct volume by 44% (p < 0.01; Fig. 2c) and improved functional outcome over 28 days (p < 0.01; Fig. 2d). In aged mice, treatment at 2 hours significantly reduced infarct volume in males (83% reduction; p < 0.05) and females (48% reduction; p < 0.01; Fig. 2c) and improved neurological outcomes over 28 days (p < 0.01; Fig. 2e,f).

In the tMCAO model, in young males, AZD1236 administered at 2 or 4, but not 6, hours significantly reduced infarct volume (25% and 22% reduction; p < 0.01 and p < 0.05, respectively Fig. 3a), and improved Garcia scores over 28 days when administered at 4 hours (p < 0.01 Fig. 3b). Longitudinal functional assessment for the 2 hours group was not available in this cohort and is therefore not shown. Similarly, in young females AZD1236 at 2 or 4 hours significantly reduced infarct volume (50% and 32% reduction; p < 0.01 and p < 0.01, respectively Fig. 3c) and improved Garcia scores (p < 0.01 Fig. 3d), In both male and female aged mice, administration at 2 hours also reduced infarct size (53% and 46% reductions; p < 0.01 and p < 0.05, respectively) and improved outcomes (p < 0.05; Fig. 3e-g). Critically, occlusion duration was adjusted to generate comparable lesion severity across biological variables, consistent with previous experimental stroke studies. Accordingly, comparisons were made within each cohort rather than interpreted as evidence of intrinsic sex-or age-dependent differences in treatment efficacy.

**Figure 3.**
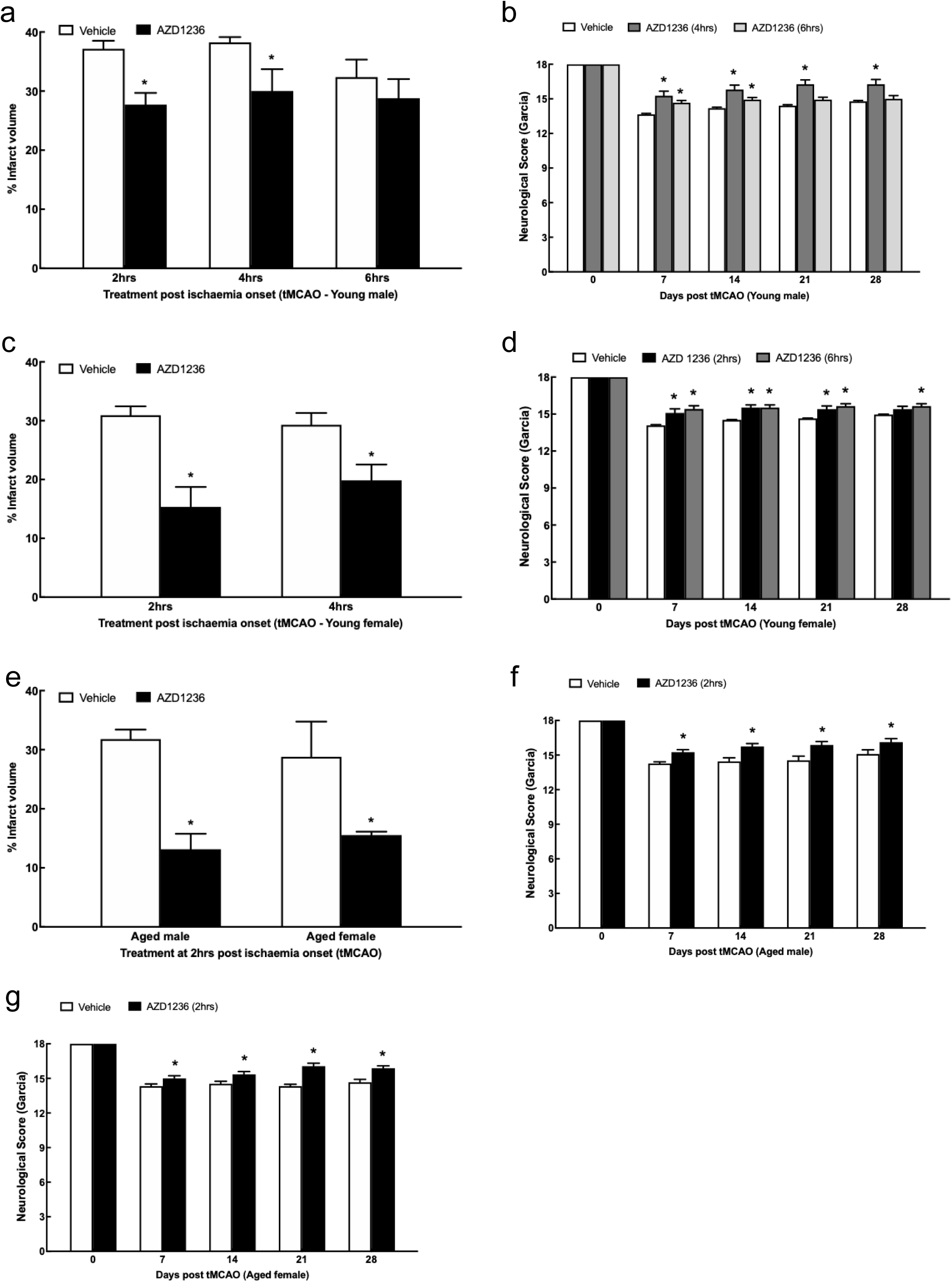
Time-dependent neuroprotection by AZD1236 following tMCAO. Infarct volume in young (a,c) and aged (e) mice, and longitudinal neurological function assessed using the 18-point Garcia score in young (b,d) and aged (f,g) mice following tMCAO. AZD1236 (200 mg/kg) or vehicle was administered 2, 4 or 6 h after ischaemia onset. Infarct-volume data were analysed using two-tailed unpaired Welch’s t tests (a,c,e) comparing each AZD1236-treated group with its contemporaneous vehicle control. Longitudinal neurological outcomes were analysed using a mixed-effects model (or two-way repeated-measures ANOVA where appropriate). *P < 0.05 versus the corresponding vehicle group.

Furthermore, in obese males with pMCAO, AZD1236 (200mg/kg) reduced infarct volume (43% reduction; p < 0.05) and improved neurological function from day 14 onwards (p < 0.01; Fig. 4a,b). Also, in obese male mice with tMCAO, AZD1236 treatment at 2 hours demonstrated a 41% reduction in infarct volume (p < 0.05) and improved neurological scores (p < 0.01; Fig. 4a,c). The magnitude of benefit in aged and obese groups was comparable to that observed in young mice.

**Figure 4.**
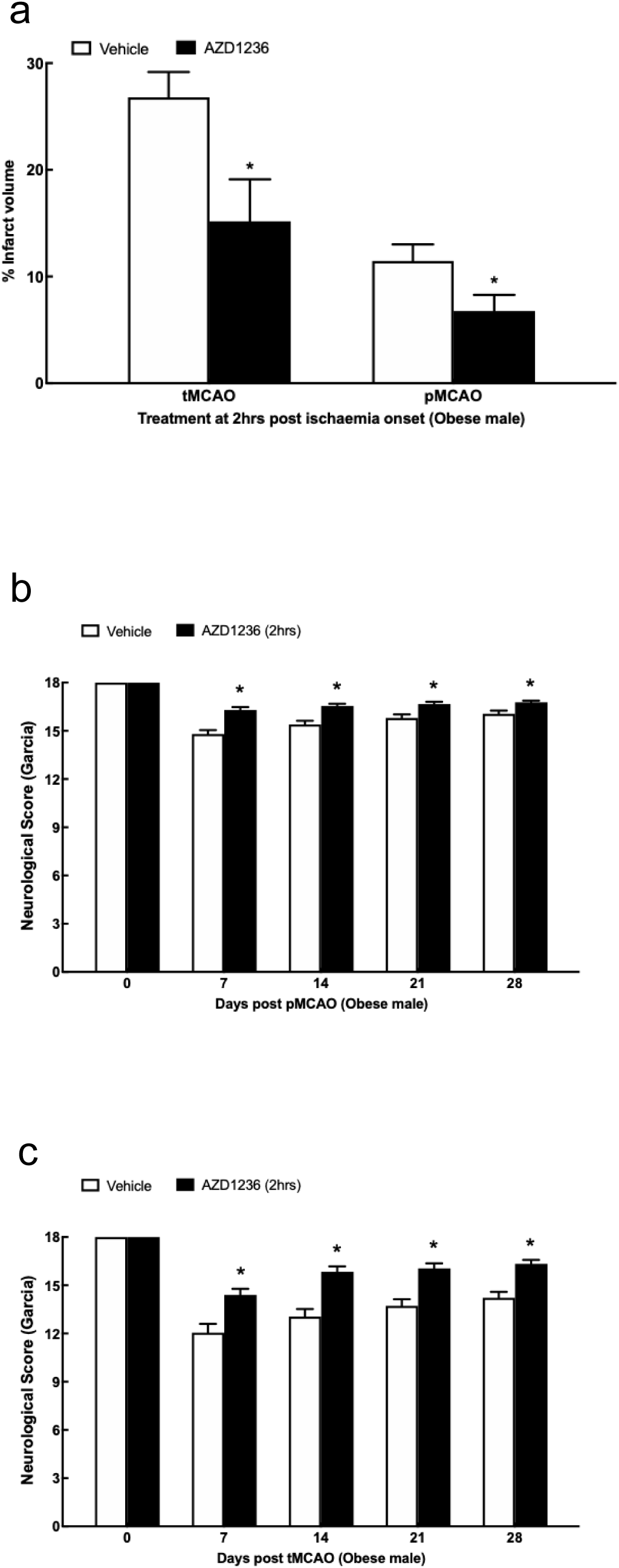
AZD1236 protects obese mice with ischaemic stroke. Obese mice received AZD1236 (200 mg/kg) or vehicle 2 h after permanent or transient middle cerebral artery occlusion. Infarct volume was assessed 48 h after stroke (a), and neurological function was assessed using the 18-point Garcia score over 28 days (b,c). Data are presented as mean ± SEM (n = 10 biological replicates per group). Infarct-volume data were analysed using two-tailed unpaired Welch’s t tests. Longitudinal neurological outcomes were analysed using a mixed-effects model (or two-way repeated-measures ANOVA where appropriate). *P < 0.05 versus vehicle.

### Effects in Haemorrhagic Stroke

In the collagenase-induced haemorrhagic stroke model, AZD1236 treatment at 2, 4, or 6 hours significantly reduced haemorrhage volume, at 48 hours, by 46%, 48%, and 67%, respectively (p < 0.01), and partially improved neurological outcome over 28 days (Fig. 5a,b).

**Figure 5.**
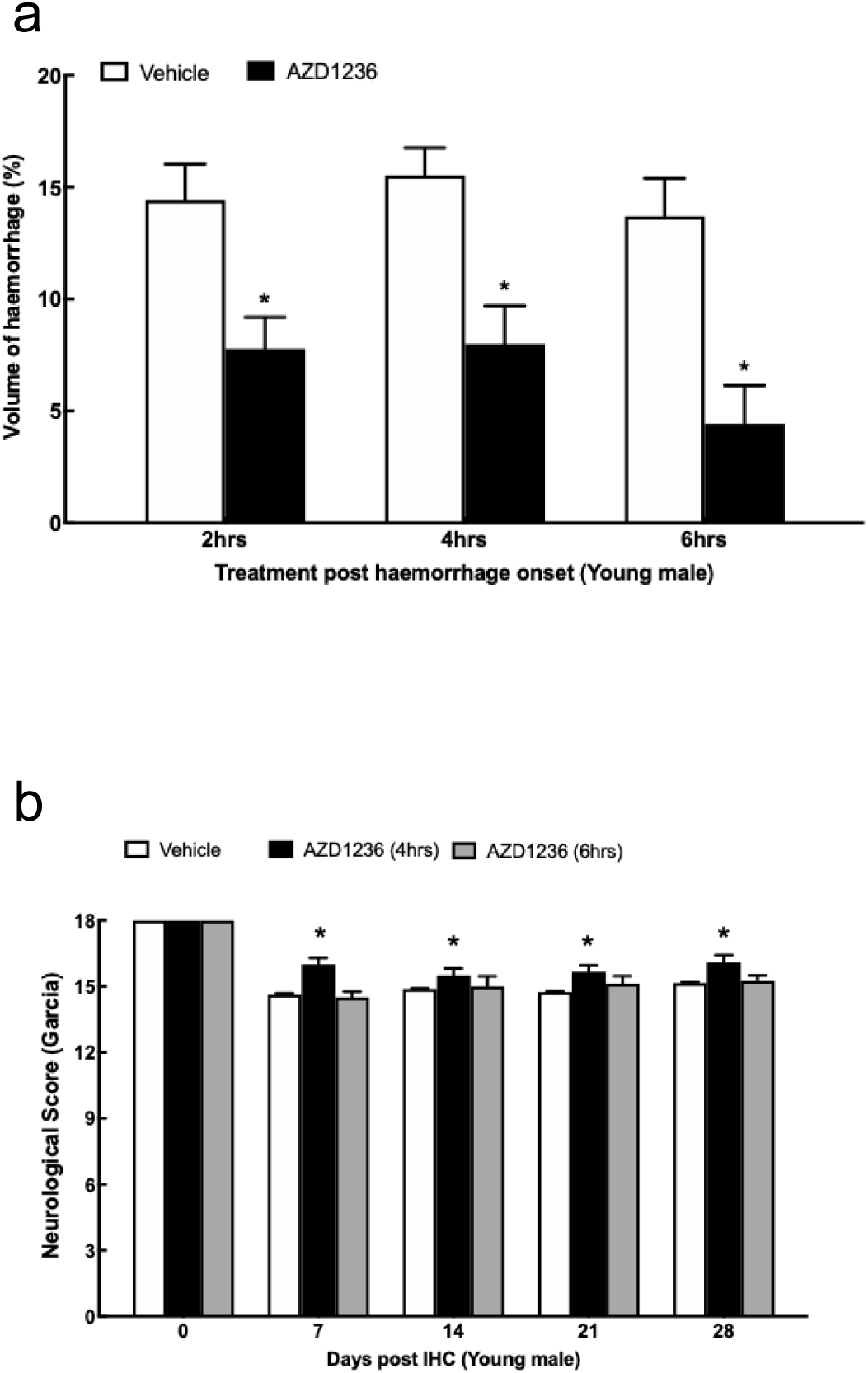
AZD1236 efficacy in haemorrhagic stroke. Haematoma volume 24 h after haemorrhage (a) and longitudinal neurological function assessed using the 18-point Garcia score over 28 days (b). AZD1236 (200 mg/kg) or vehicle was administered 2, 4 or 6 h after haemorrhage induction. Data are presented as mean ± SEM (n = 8–13 biological replicates per group). Haematoma-volume data were analysed using one-way ANOVA followed by Dunnett’s multiple-comparisons test. Garcia scores were analysed using a mixed-effects model (or two-way repeated-measures ANOVA where appropriate). *P < 0.05 versus vehicle.

### Preservation of Blood-Brain Barrier integrity

AZD1236 significantly reduced BBB leakage following ischaemia. In the pMCAO model, treatment at 2 hours reduced Evans Blue extravasation in brain and spinal cord tissue (Fig. 6a). In the tMCAO model, a significant reduction in Evans Blue accumulation was observed in brain tissue (Fig. 6b). MRI studies confirmed improved BBB integrity in treated mice: gadolinium enhancement was greater in vehicle-treated than AZD1236-treated animals, and the reduction in BBB permeability with AZD1236 was statistically significant (p < 0.01; Fig. 7). Representative pre-and post-contrast MRI images and derived maps are provided in Supplementary Fig 12.

**Figure 6.**
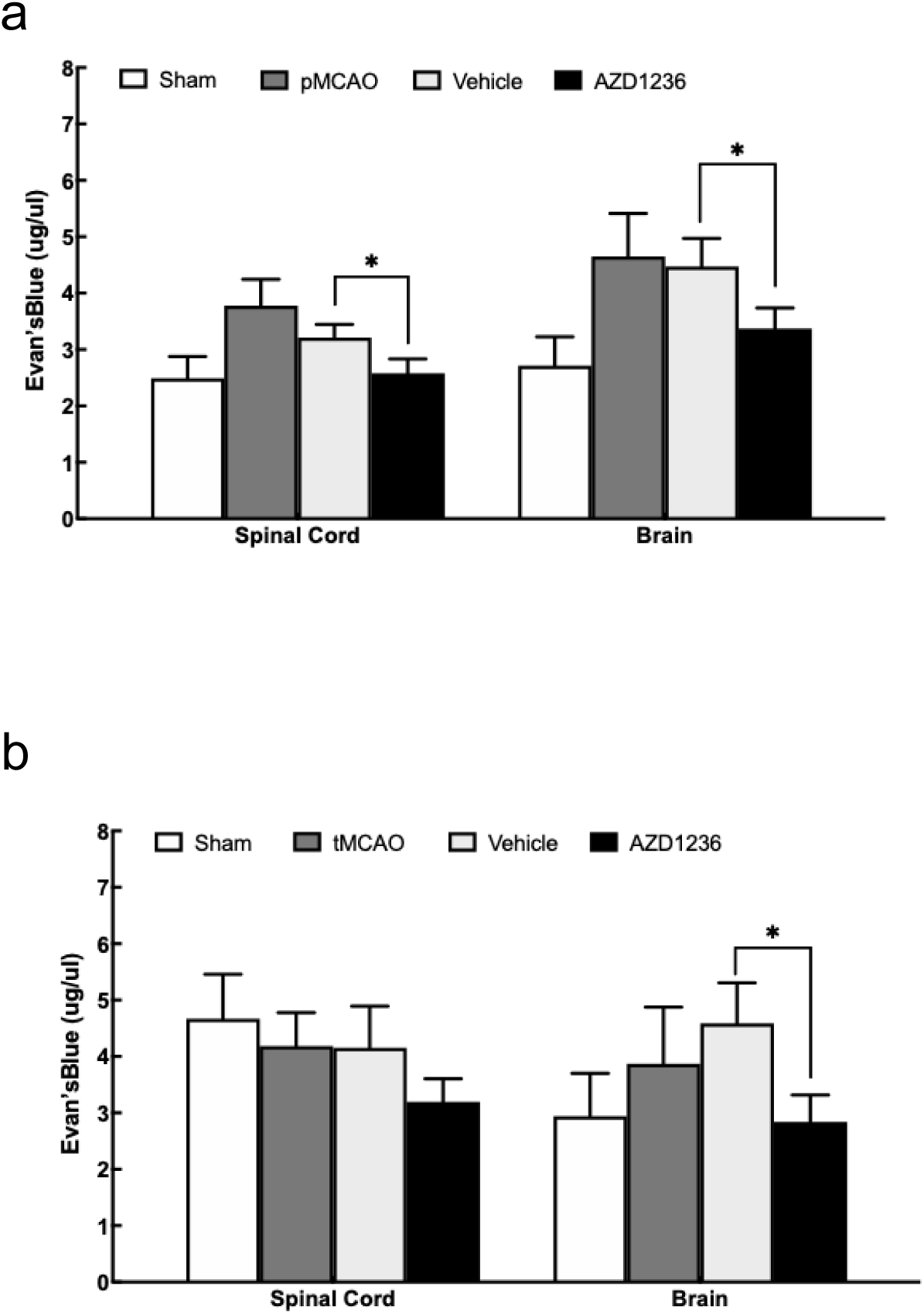
AZD1236 reduces blood–brain barrier breakdown after ischaemia. Evans Blue extravasation was measured 48 h after permanent (a) or transient (b) middle cerebral artery occlusion in young male mice treated with AZD1236 (200 mg/kg) or vehicle 2 h after ischaemia. Groups comprised sham, stroke alone, stroke + vehicle and stroke + AZD1236. Data are presented as mean ± SEM (n = 8 biological replicates per group) and were analysed using one-way ANOVA followed by Dunnett’s multiple-comparisons test. *P < 0.05 versus stroke + vehicle.

**Figure 7.**
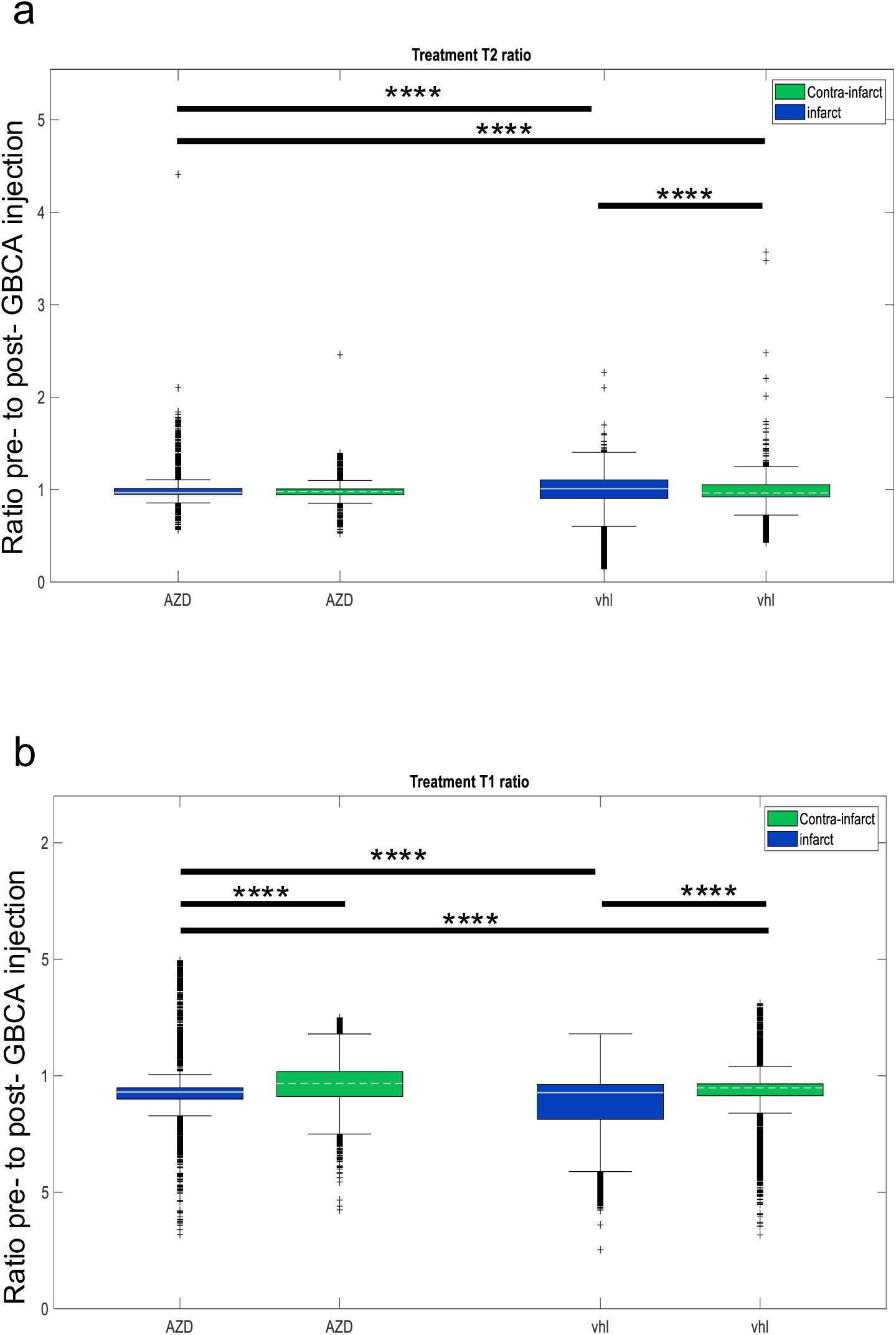
MRI evidence of improved blood–brain barrier integrity with AZD1236 after tMCAO. Quantitative MRI assessment of blood–brain barrier permeability 24 h after tMCAO in young male mice treated with vehicle or AZD1236 (200 mg/kg) 2 h after ischaemia. Ratios of pre-to post-gadolinium signal intensity were quantified in cortical and subcortical regions. Data are presented as box-and-whisker plots showing the median, interquartile range (IQR), and 1.5 × IQR. Outliers are indicated by (+). n = 8 biological replicates per group. Statistical analysis was performed as described in the Methods. ****P < 0.0001.

### Molecular mechanisms

Western blot analysis at day 3 post-ischaemia showed that AZD1236 significantly reduced both MMP-9 and MMP-12 expression compared with vehicle controls (p < 0.01; Fig. 8a,b). Treatment also increased the expression of the tight junction proteins claudin-5 and ZO-1 (p < 0.05; Fig. 8a,c), consistent with preservation of BBB integrity. Levels of phospho-ERK1/2 and cleaved caspase-3 were unaffected (data not shown), suggesting that the drug’s effects are not mediated through apoptotic or ERK-dependent signalling pathways.

**Figure 8.**
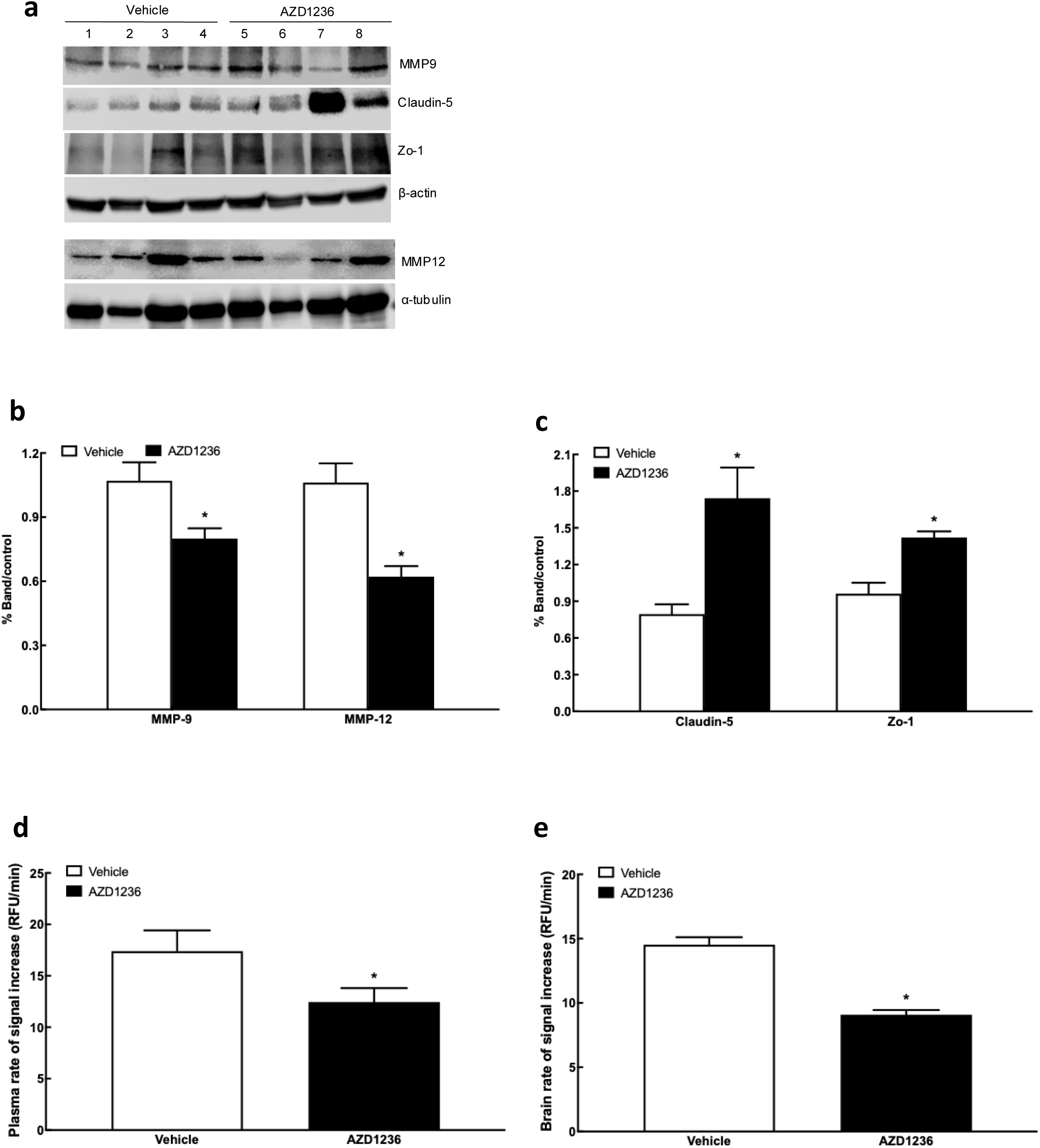
AZD1236 enhances tight junction protein expression and brain and plasma gelatinase inhibition following ischaemia. Representative immunoblots (a) and quantification of MMP-9 and MMP-12 expression (b), together with quantification of the tight junction proteins claudin-5 and zona occludens-1 (c), in brain tissue 3 days after tMCAO. Gelatinase inhibitory capacity was assessed in plasma (d) and brain (e) 24 h after tMCAO using the DQ-gelatin assay. Data are presented as mean ± SEM (n = 4–5 biological replicates per group) and were analysed using two-tailed unpaired Welch’s t tests. *P < 0.05 versus vehicle.

### AZD1236 inhibits gelatinase activity following pMCAO

To determine whether AZD1236 treatment resulted in sustained metalloproteinase inhibition, circulating gelatinase inhibitory capacity was assessed in plasma and brain at 24 hours post-pMCAO using a fluorogenic DQ-gelatin assay. Both plasma and brain from vehicle-treated mice exhibited measurable gelatinase activity when incubated with Clostridium collagenase, whereas plasma and brain from AZD1236-treated mice showed significantly reduced substrate cleavage, indicating increased inhibitory capacity (Fig. 8c,d). Quantification of fluorescence kinetics demonstrated a significant reduction in gelatinase activity (ΔRFU/min) in the AZD1236-treated group compared with vehicle controls (p < 0.05). These data indicate that early AZD1236 administration (2 hours after ischaemia onset) produces sustained circulating gelatinase inhibitory activity detectable at 24 hours following ischaemic stroke.

### Post-stroke pain behaviour

In the pMCAO model, young male mice treated at 2 or 4 hours exhibited reduced mechanical allodynia (von Frey test; Fig. 9a) and thermal hyperalgesia (Hargreaves test; Fig. 9c) compared with vehicle controls (p < 0.01). No analgesic benefit was observed when treatment was initiated at 6 hours. Similar reductions in hypersensitivity were observed in young female mice treated at 2 or 4 hours (Fig. 9b,d). These improvements occurred without changes in cleaved caspase-3, suggesting that analgesic effects are linked to BBB preservation and suppression of MMP-9 and MMP-12 activity rather than reduced apoptosis. Furthermore, these responses cannot be attributed to impaired motor function following stroke: animals retained withdrawal reflexes in response to mechanical stimulation, indicating that the underlying sensorimotor circuitry remained intact. Therefore, the changes observed, particularly in vehicle-treated animals, are unlikely to reflect an inability to move the affected hindlimb but rather altered sensory processing and nociceptive sensitivity.

**Figure 9.**
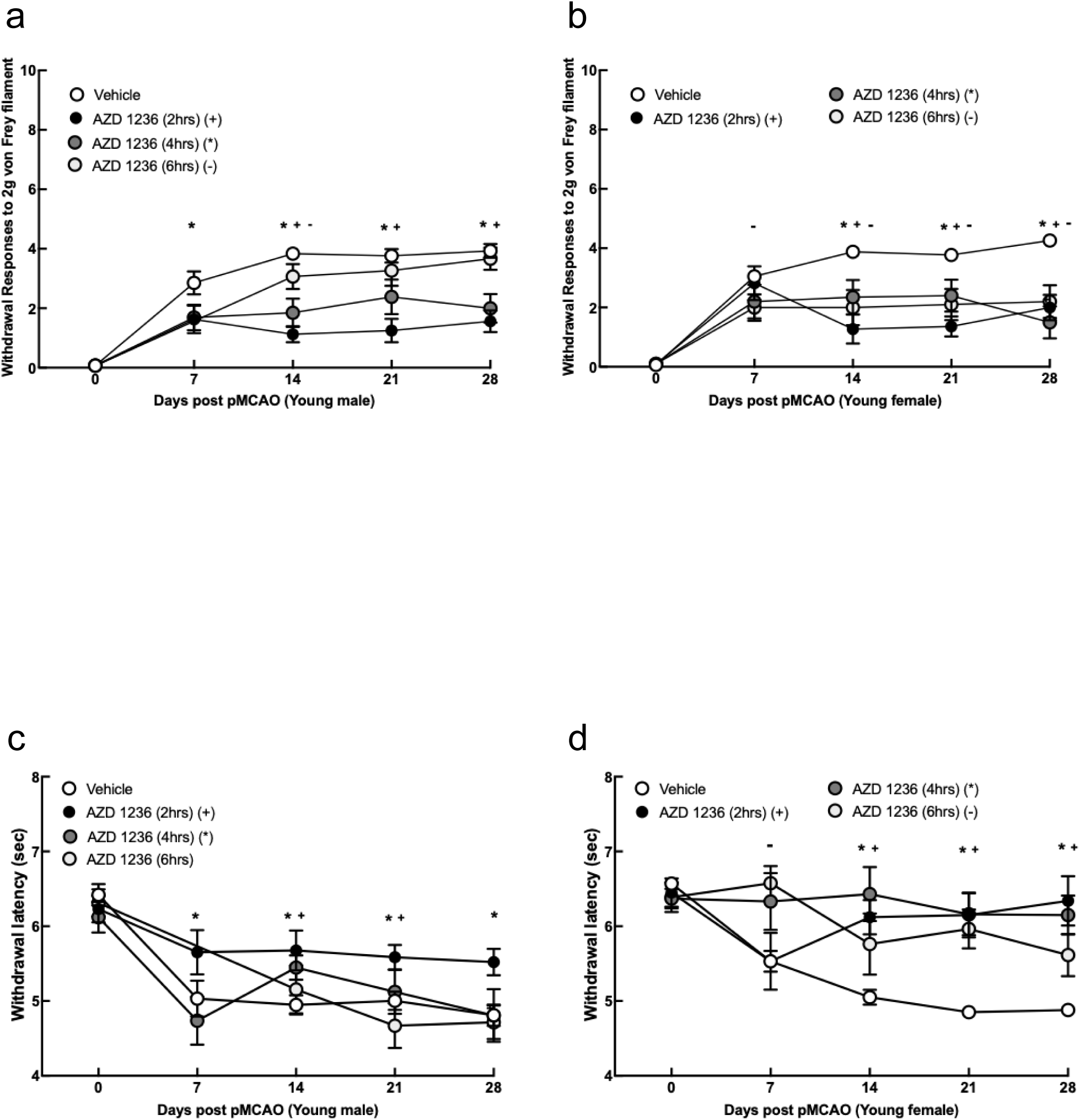
AZD1236 reduces development of hypersensitivity following ischaemic stroke. Mechanical allodynia (a,b) and thermal hyperalgesia (c,d) were assessed over 28 days after pMCAO in young male and female mice treated with AZD1236 (200 mg/kg) or vehicle 2, 4 or 6 h after ischaemia, followed by once-daily dosing for 7 days. Data are presented as mean ± SEM (n = 10–15 biological replicates per group). Longitudinal behavioural data were analysed using a mixed-effects model (or two-way repeated-measures ANOVA where appropriate). P < 0.05 versus the corresponding vehicle group. Symbols denote treatment initiated at 2 h (*), 4 h (+) and 6 h (−) after stroke.

## DISCUSSION

This study demonstrates that the dual MMP-9/MMP-12 inhibitor AZD1236 provides robust neurovascular protection across multiple experimental stroke paradigms. Treatment initiated within 2 - 4 hours of stroke reduced infarct volume, preserved blood–brain barrier integrity, improved long-term neurological outcome, and attenuated the development of post-stroke pain. Importantly, efficacy was observed in permanent and transient ischaemia, intracerebral haemorrhage, both sexes, aged animals, and animals with obesity, supporting the robustness and potential translational relevance of these findings (11,15, 16).

### Key Findings

The consistency of benefit across multiple models and outcome measures supports a central role for MMP inhibition in limiting secondary neurovascular injury after stroke. The therapeutic window of 2 - 4 hours corresponds closely with the early post-ischaemic upregulation of MMP-9 and MMP-12, while the loss of efficacy at 6 hours suggests that intervention must occur during the initial phase of BBB disruption and inflammatory activation. In addition, AZD1236 reduced haematoma volume and improved outcome in a collagenase-induced intracerebral haemorrhage model, suggesting broader neurovascular protection across both ischaemic and haemorrhagic stroke. Furthermore, the observed efficacy of AZD1236 administered at 6 hours post haemorrhage, may be due to secondary injury linked to MMP activation that drives the haematoma expansion for longer, which is not observed in the ischaemic model. Therefore, later inhibition remains beneficial.

Mechanistically, AZD1236 reduced expression of MMP-9 and MMP-12 and upregulation of the tight junction proteins claudin-5 and ZO-1 supporting a mechanism of BBB preservation. It is likely that the observed neuroprotection arises from the combined suppression of MMP-9 and MMP-12 activity rather than reduced expression alone. This dual inhibition is mechanistically attractive, as MMP-9 drives early BBB disruption (28, 31, 74), while MMP-12 amplifies inflammation and matrix degradation in a delayed phase (31–33, 64). Gelatinase activity following ischaemic stroke is most commonly attributed to MMP-9, with its central role in BBB disruption, neuroinflammation, and extracellular matrix remodelling. However, AZD1236 is a dual inhibitor targeting both MMP-9 and MMP-12. While MMP-9 is likely the primary contributor to the gelatinolytic signal measured in the DQ-gelatin assay, MMP-12 (macrophage elastase) is also implicated in post-ischaemic inflammatory responses, including leukocyte infiltration and secondary tissue injury. Therefore, the effects observed here are likely mediated by the combined inhibition of both MMP-9 and MMP-12 rather than a single protease target. This broader inhibition profile may be advantageous in modulating the complex proteolytic and inflammatory environment following stroke. The parallel reduction in gelatinase activity observed in both plasma and brain suggests that AZD1236 exerts both systemic and central effects and raises the possibility that peripheral MMP inhibition may contribute to central outcomes, either through direct penetration or through modulation of circulating inflammatory mediators that influence the neurovascular unit.

Our findings also demonstrate that AZD1236 preserves BBB function, as evidenced by reduced Evans Blue extravasation, lower gadolinium leakage on MRI, and increased expression of tight junction proteins. BBB dysfunction is a recognised driver of neuroinflammation, secondary neuronal injury, haemorrhagic transformation, and chronic neurological decline after stroke (17, 18, 22, 25, 27). Pharmacological preservation of the BBB is therefore likely to contribute to the long-term functional recovery observed in this study. The limited penetration of AZD1236 across an intact BBB, while initially a potential limitation, may represent an advantage: the compound preferentially enters the injured brain during the acute pathological phase, reducing the risk of interfering with later reparative processes such as angiogenesis and remodelling (39, 43, 74, 75). This profile aligns with the concept of a targeted neurovascular therapy that acts during the window of maximal BBB disruption and MMP activation.

A novel finding of this study is that AZD1236 reduces mechanical allodynia and thermal hyperalgesia when administered 2-4 hours after stroke. Pain phenotyping was restricted to young adult mice to reduce complications due to age-related motor decline. Central post-stroke pain is a debilitating complication affecting a substantial proportion of survivors and is strongly linked to neuroinflammatory signalling (48–50, 58). The present data suggest that dual inhibition of MMP-9 and MMP-12, with consequent attenuation of BBB disruption and inflammatory cascades, may interrupt the pathways that lead to central sensitisation. The ability of a single agent to provide both acute neuroprotection and long-term mitigation of post-stroke pain represents a potentially important therapeutic advance. Notably, stroke did not abolish the withdrawal reflex, supporting the conclusion that behavioural changes reflect altered nociceptive processing rather than motor impairment, thereby strengthening the interpretation of the pain-related outcomes. These findings are consistent with previous work implicating MMPs in neuropathic pain and extend this concept to central post-stroke pain states.

### Translational Implications and Limitations

The broad efficacy of AZD1236 across experimental models, sexes, and clinically relevant comorbid conditions supports its translational potential. The 2–4-hour therapeutic window aligns with the time frame in which many patients currently present to hospital, suggesting that AZD1236 could be deployed either alone or in combination with reperfusion therapies. AZD1236 has been administered to healthy volunteers at single doses of up to 1,500 mg and repeat doses of up to 500 mg daily for 13 days, and at 75 mg twice daily for six weeks in moderate-to-severe COPD, with acceptable tolerability (45, 46). Allometric scaling of 200 mg/kg gives a human-equivalent dose of approximately 16mg/kg (approximately 1,100 mg for a 70 kg adult) daily. However, such scaling does not account for interspecies differences in pharmacokinetics, protein binding or target potency, for examplebecause the human enzymes are 20-to 50-fold more sensitive, comparable target coverage would be anticipated at considerably lower exposures. Accordingly, the present study should be viewed primarily as a proof-of-pharmacology investigation rather than a dose-optimisation study.

Nonetheless, several limitations warrant consideration. Daily dosing for 7 days was evaluated; further dose–response and pharmacokinetic studies may help to refine optimal regimens and define minimally effective exposures. While mouse models provide essential proof-of-concept, they cannot fully replicate the complexity of human stroke, including comorbidities, heterogeneous lesion patterns, and variable reperfusion. These findings should therefore be interpreted primarily as evidence of pharmacodynamic target engagement and neurovascular protection rather than definitive proof of clinical efficacy, and further studies will be required to establish causal links between specific MMP inhibition profiles and behavioural outcomes.

Furthermore, the fluorogenic gelatinase assay does not discriminate between individual MMP species and therefore does not allow direct attribution of the observed effects to specific proteases. However, interpretation can be supported by the known selectivity profile of AZD1236, which preferentially targets MMP-9 and MMP-12, together with the established role of MMP-9 as a major contributor to gelatinolytic activity following ischaemic stroke. As such, the observed reduction in substrate cleavage is most consistent with inhibition of MMP-9-dominated activity, while also reflecting potential contributions from MMP-12 and other gelatinases within the complex post-ischaemic proteolytic environment. Finally, the precise mechanisms underlying the analgesic effects require further investigation, particularly in relation to microglial activation, synaptic plasticity, and downstream inflammatory mediators.

### Conclusions

This study provides compelling preclinical evidence that dual MMP-9/MMP-12 inhibition with AZD1236 reduces infarct volume, preserves BBB integrity, and improves long-term neurological function across clinically relevant stroke models and populations. In addition, AZD1236 attenuates the development of central post-stroke pain, broadening its therapeutic scope beyond the acute phase. These findings support AZD1236 as a strong candidate for further translational development and future clinical trials aimed at addressing the significant treatment gap in stroke management.

The choice of 200 mg/kg was intended as a high-dose pharmacologic challenge to drive near-maximal MMP-9/12 inhibition rather than to define a minimally effective dose. Future studies should explore lower doses and alternative dosing schedules to determine the minimal exposure required for robust neurovascular protection and to facilitate translation to clinically achievable regimens.

## Acknowledgement

The authors gratefully acknowledge the staff of University of Sheffield Statistical Services Unit, in particular Pete J Laude for his invaluable statistical advice and support throughout the project and during manuscript preparation. The authors would like to acknowledge the staff of the University of Sheffield Biological Services Unit for their excellent animal care and support.

**Supplemental material Figure 10 PK in Obese mice.** Brain and plasma concentrations of AZD1236 following tMCAO in obese male mice. AZD1236 (100 mg/kg) was administered 2 h after ischaemia onset and concentrations were measured over 24 h. Data are mean ± SEM (n = 5–6 biological replicates per group).

**Supplemental material Figure 11 Histology images.** Representative TTC-stained brain sections illustrating infarct size following pMCAO in young male (a), young female and aged female (b), and tMCAO in young male (c), young female (d) and aged female (e) mice. Representative haemorrhage images from the collagenase-induced intracerebral haemorrhage model (f).

**Supplemental material Figure 12 MRI map**: Representative pre-contrast T1 (a) and T2 (b) maps and post-contrast T1 (c) and T2 (d) maps. Structural images (top row) indicate regions of interest (ROIs) corresponding to the infarct, contralateral hemisphere and whole brain. The second row shows quantitative T1/T2 maps, the third row shows the corresponding goodness-of-fit (R²) maps, and the fourth row shows ROI histograms.

## Supplemental methods

### LC-MS Quantitative Analysis of AZD1236 in Serum and Brain Samples

Preparation of standards

A stock solution of AZD1326 was created by adding the appropriate amount of drug to a volume of 0.2% formic acid in acetonitrile to a final concentration of 10 mM. The solution was then sonicated at 30 ◦C for 10 minutes. This stock solution was then used to create standards at the following concentrations: 500000, 50000, 30000, 15000, 5000, 1500, 500, 150 and 50nM. This process was repeated for the internal standard, AZD3342.

### LC-ESI-MRM-MS/MS

Normal phase liquid chromatography was performed using an Agilent Infinity II 1290 high performance liquid chromatography (HPLC) system. Solvent A (5% Acetonitrile, 95% HPLC grade water with 0.1% formic acid) and solvent B (95% Acetonitrile, 5% HPLC grade water with 0.1% formic acid) were used along with a C18, 5µ, 40 x 2.1 mm (Waters Ltd, UK), with the column temperature set to 30°C. Peptides were eluted at 0.4 mL/min with a gradient elution profile from 100% A to 100% B over 13 minutes. An Agilent Ultivo triple quadrupole mass spectrometer was used with Agilent Jet Stream Electrospray Ionisation (AJS-ESI) source. Electrospray ionization was performed in positive polarity mode, with source parameters as follows: sheath gas temperature = 250°C; sheath gas flow rate = 11 L/min; desolvation gas temperature = 300°C; desolvation gas flow rate = 7 L/min; nebuliser gas pressure = 15 psi; capillary voltage = 4000 V; nozzle voltage = 1500 V. The mass spectrometer was operated in a dynamic multiple reaction monitoring (dMRM) mode using as collision energy of 33eV. Data acquisition and analysis were achieved using MassHunter Acquisition and Qualitative Analysis version B.0.10.0. The following ion transitions were used AZD1236 (m/z 416.2 to 197.1) and AZD3342 (m/z 402.2 - 190.2).

### Method Optimisation

-LC-MS

Fragmentor voltage and collision energy were optimised for each MRM target to increase assay sensitivity. 10 µL of each standard was injected in triplicate at a range of fragmentor voltages in 10 V steps, and at the default voltage of 135 V. Once the fragmentor voltage was optimised for each compound, the method was adjusted accordingly, and the collision energy was optimised for each transition. 10 µL of each standard was injected in triplicate at a range of collision energies from 0 to 40 V.

-Standard Curves and Quality Control

Serial dilutions of the standard (AZD3342) were prepared to create a series of 19 standards ranging from 0.05 to 500 µM. Standards were injected in a randomised order at the beginning and end of each experimental run, and after analysis of each sample set. 10 µL of blank sample containing 50% acetonitrile with 0.1% formic acid was injected at the beginning and end of each experimental run, before and after each set of standards, and after every 10 sample injections.

### MRI data analysis

Custom Matlab code (MathWorks, Natick, MA) was used to create T1 and T2 maps of each brain slice. For each voxel position the image intensity was fitted by a non-linear least squares algorithm to either:

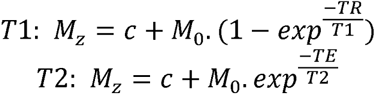

Where Mz is the signal intensity; M0 the equilibrium magnetisation; TR the acquisition repetition time; TE the echo time and c is an offset the aid fitting optimisation.

After T1/T2 map generation a region of interest (ROI) was drawn around the whole brain and stroke infarct area (this was guided by both a brighter brain contrast in the magnitude image and elevated T2 values in the T2 map. A contra-lateral region to the stroke area was created by reflecting the infarct ROI about the brain centreline for each slice (contra-infarct). T1/T2 value were taken for slices containing infarct and contra-infarct regions for statical analysis. Representative images and T1/T2 maps are shown in Figure S1/2.

To analyse the effect of Gadolinium (Gd) on tissue perfusion, the ratio of pre and post IP Gd injection T1 (or T2) values from the infarct and contra-infarct regions was determined. As the animal had been moved to perform the injection the image slices were not from exactly the same region. Additionally, due to a change in contrast, the total number of ROI voxels was different between pre and post injection. To account for this the number of voxels (ROI*slices) from the smaller of the pre and post Gd data sets was found and the same number of voxels randomly selected from the larger data set. For both pre and post injection T1 and T2 values for each ROI were numerically sorted in ascending order and divided per voxel (i.e. Post Gd voxel 1/Pre Gd voxel 1, Post Gd voxel 2/Pre Gd voxel 2). A ratio of <1 indicates that the T1/T2 had decreased post Gd. Once this analysis was completed for all mice, the treatment was then unblinded and T1/T2 data collected into infarct/contra-infarct and AZD/vehicle groups. Using Matlab, statistical analysis was by Kruskal-Wallis with a Bonferroni post-hoc test.

