## Supplementary figures and images for "Dual MMP-9/12 Inhibition with AZD1236 Confers Neurovascular Protection and Reduces Post-Stroke Pain in Experimental Stroke Models"

### Supplementary Fig 10

#### Slide 1
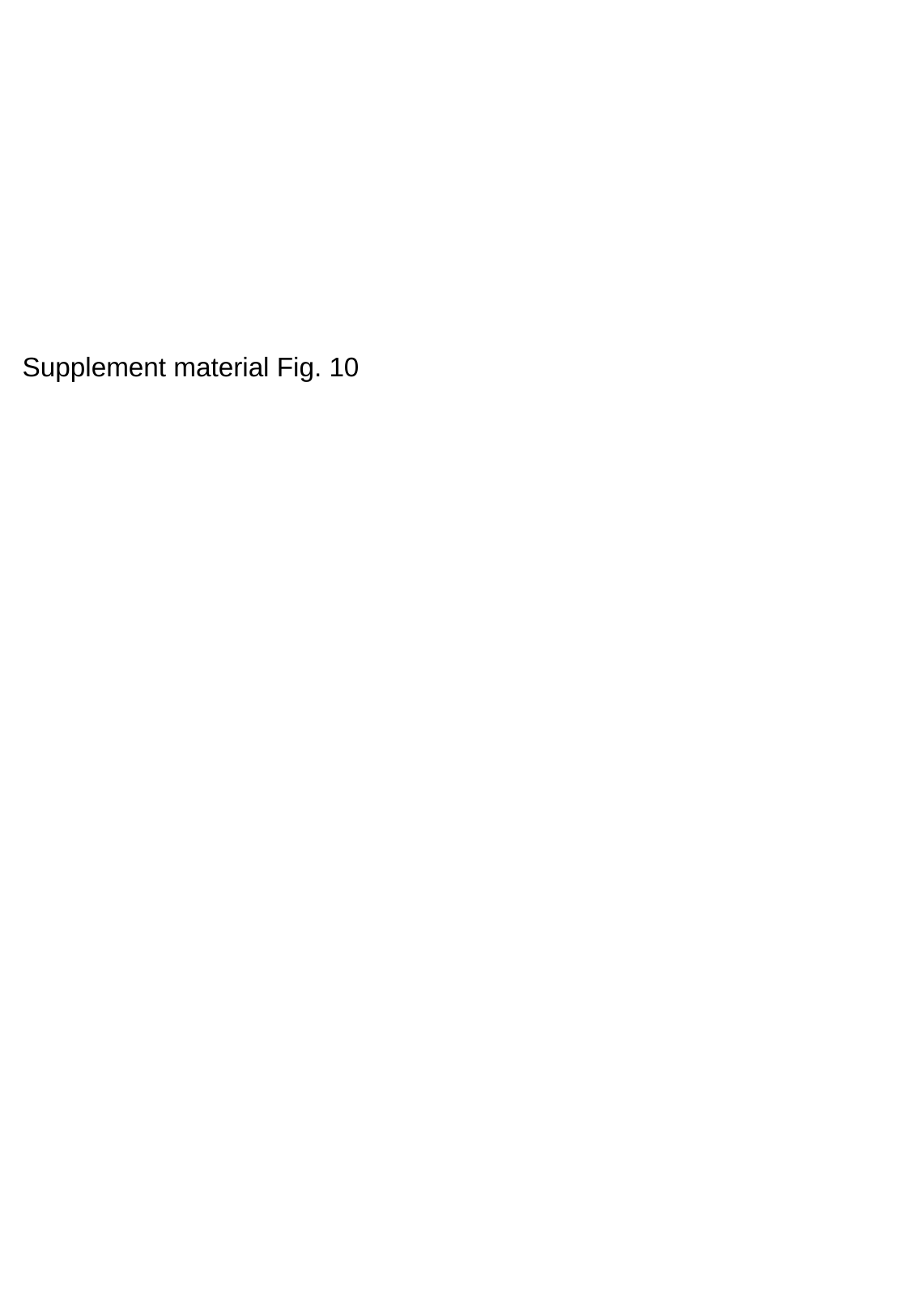

Supplement material Fig. 10

### Supplementary Fig 12

#### Slide 1
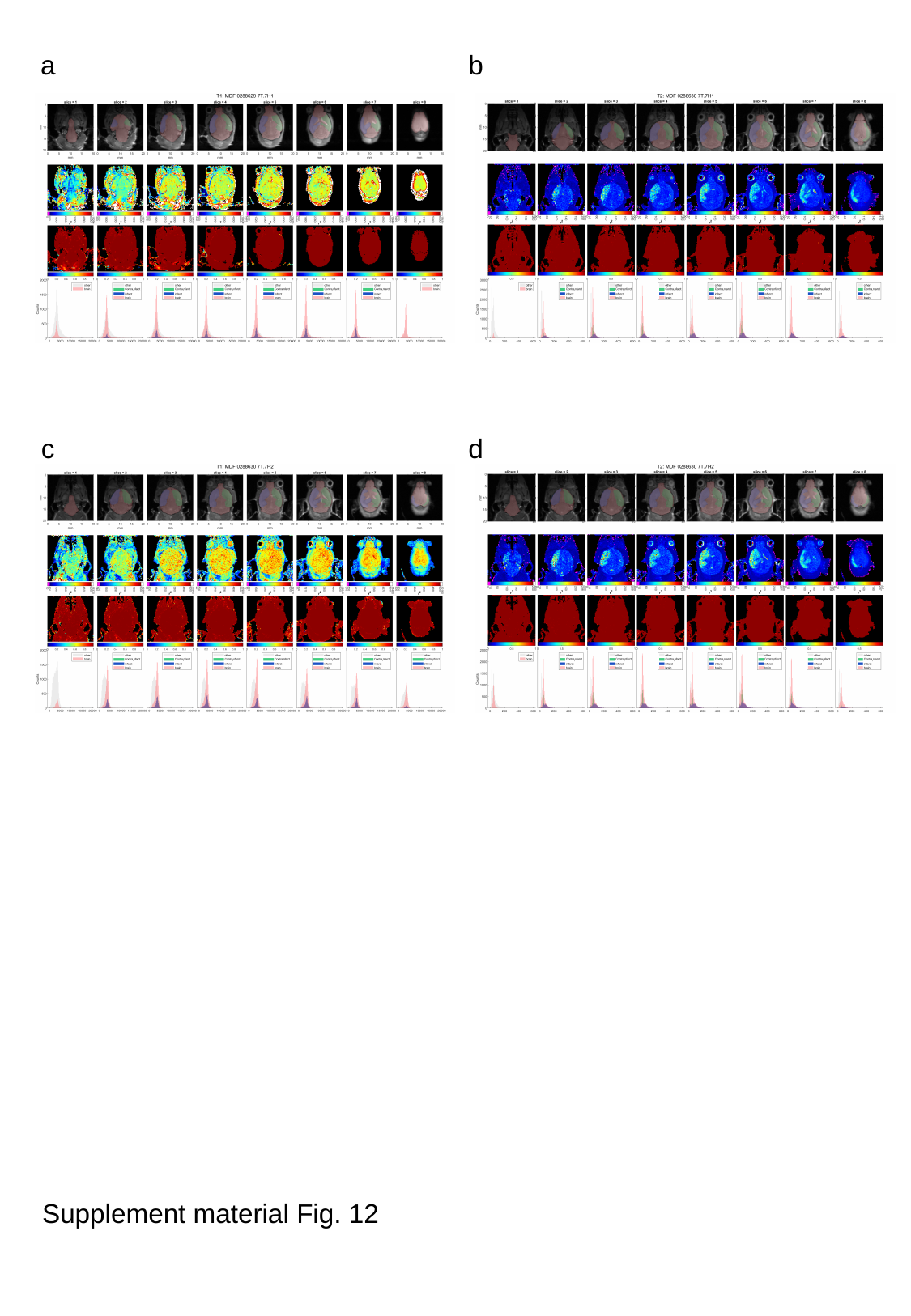

a
b
c
d
Supplement material Fig. 12
