## Supplementary Fig 11 for "Dual MMP-9/12 Inhibition with AZD1236 Confers Neurovascular Protection and Reduces Post-Stroke Pain in Experimental Stroke Models"

### Slide 1
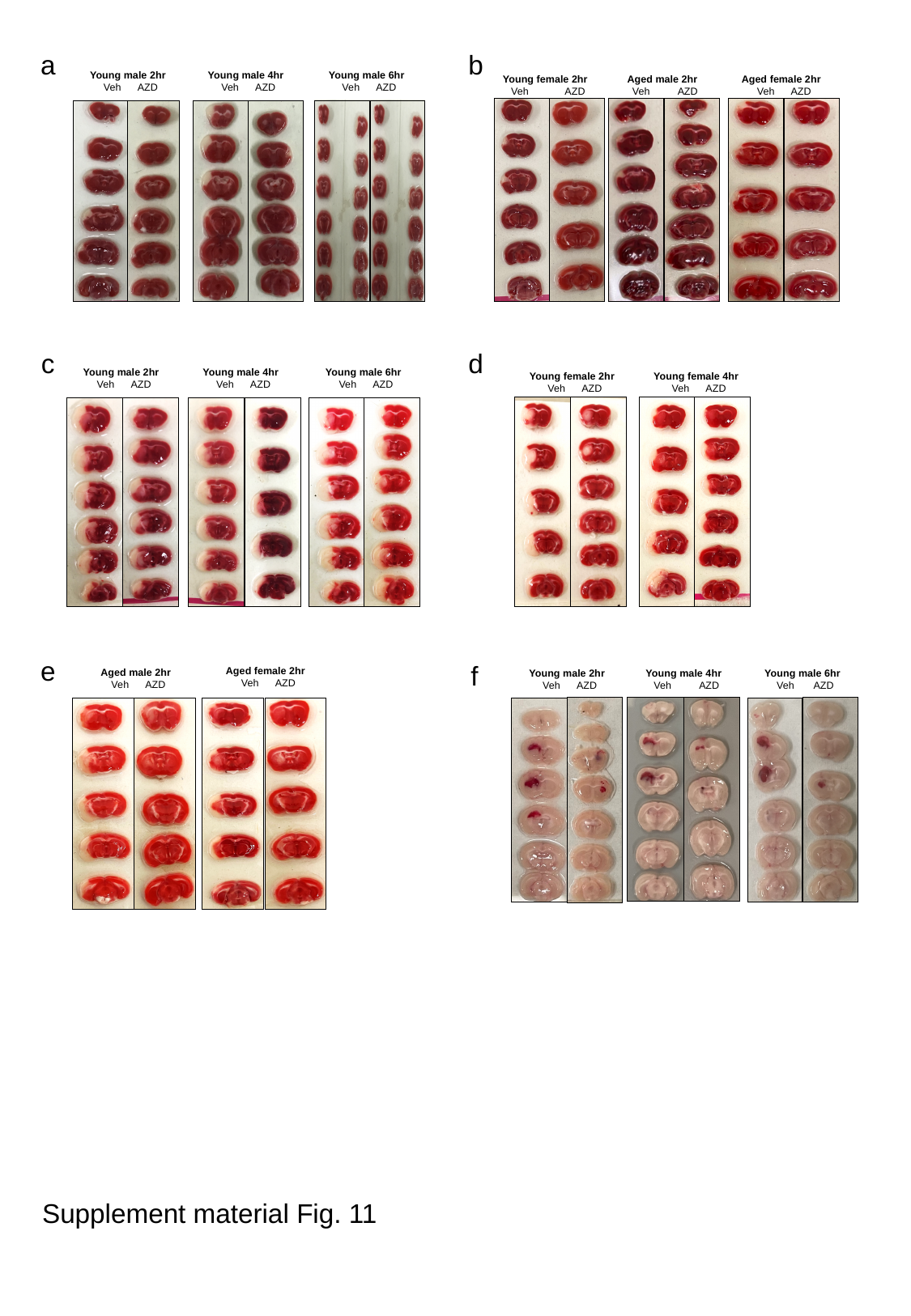

a
b
Young male 2hr
 Veh AZD
Young male 4hr
 Veh AZD
Young male 6hr
 Veh AZD
Young female 2hr
 Veh AZD
Aged male 2hr
 Veh AZD
Aged female 2hr
 Veh AZD
c
d
Young male 2hr
 Veh AZD
Young male 4hr
 Veh AZD
Young male 6hr
 Veh AZD
Young female 2hr
 Veh AZD
Young female 4hr
 Veh AZD
e
Aged female 2hr
 Veh AZD
Aged male 2hr
 Veh AZD
f
Young male 2hr
 Veh AZD
Young male 4hr
 Veh AZD
Young male 6hr
 Veh AZD
Supplement material Fig. 11
